# HeartVar: An LLM-Assisted Tool for Clinical Classification of Variants in Cardiovascular Disease Cohorts

**DOI:** 10.64898/2026.09.10.750569

**Authors:** Jamie-Lee Thompson, Debjani Das, Sally L. Dunwoodie, Eleni Giannoulatou

## Abstract

Manual clinical DNA variant classification is the bottleneck of every clinical and research rare disease workflow. The process typically requires a curator to assemble evidence from numerous databases, weigh 28 criteria, reconcile competing evidence, and produce a defensible case for the final classification. Additionally, the framework used to assess variants is not static and successive addenda have revised individual criteria. The most complete and current evidence aggregators available are commercial platforms, which can limit researcher access. We present HeartVar, an open-source web tool that automates the evidence-gathering and interpretation steps of variant classification associated with cardiovascular disease. Given a gene, a variant, and clinical context, HeartVar queries 20 public databases in parallel and assigns ACMG/AMP criteria through a hybrid rule-based/large language model (LLM) approach. Criteria that can be resolved from structured data are computed programmatically, and only those requiring interpretation of unstructured evidence are passed to the LLM. HeartVar returns a classification, point score, per-criterion breakdown, clinical-narrative summary, and database annotations. Benchmarking of 106 expert-curated ClinGen variants showed HeartVar outperformed other curation tools, assigning the correct ACMG tier in 72% of cases. HeartVar demonstrates that an LLM constrained by a domain-specific prompt and grounded in structured evidence can produce variant interpretations of first-pass quality for a cardiovascular disease cohort; however, it is not intended to replace manual assessment by a qualified variant curator. The tool is freely available to use and hosted at www.heartvar.victorchang.edu.au.

## Introduction

Clinical classification of DNA variants according to the ACMG/AMP framework is the standard for clinical and research reporting of germline sequence variation (1). The framework specifies 28 evidence criteria across which a curator must evaluate, weigh according to their strengths, and combine into one of five classification categories: Pathogenic (P), Likely Pathogenic (LP), Variant of Uncertain Significance (VUS), Likely Benign (LB), or Benign (B). The Tavtigian et al. (2020) point-based system formalised the process of combining numerical value points of each evidence strength (2), and a continuing series of ClinGen Sequence Variant Interpretation (SVI) addenda have refined the strength of individual criteria for specific data types (3–6).

Applying the framework to a single variant requires evidence drawn from a heterogeneous set of databases including population frequency (gnomAD) (7), in-silico prediction (CADD, SIFT, PolyPhen, REVEL, AlphaMissense, SpliceAI) (8–13), prior clinical reports (ClinVar) (14), gene-disease validity (GenCC, PanelApp) (15, 16), protein context (UniProt, ProtVar) (17, 18), expression (GTEx) (19), functional support (MGI) (20), literature (PubMed), and interactions (BioGRID) (21). Each differs in structure, terminology, and query interface meaning an experienced curator can spend hours gathering evidence for each variant. Further, expert curators routinely disagree on criteria selection and the strength at which they apply (22).

Software support for variant classification has developed along four largely separate lines (Table 1). Rule-based automation such as InterVar (23), TAPES (24), AutoGVP (25), BIAS-2015 (26), and ACVI-Med (27), annotate a variant and evaluate the programmatically. These tools are open, auditable, and free, but are disease-agnostic and largely unable to apply criteria that require judgement (e.g. PS3/BS3, PP4). GeneBe (28) evaluates 17 of the 28 criteria and performs comparably to commercial platforms, but is closed source and heavily reliant on ClinVar evidence. Disease-specific variant curation tools, such as CardioClassifier (29) and CardioVAI (30), showed that disease-specific parameterisation identifies more clinically actionable variants with fewer false positives. However, neither tool released their source code, leaving users unable to update, audit, or adapt them; CardioClassifier has not been updated since 2020, and CardioVAI’s free server has been absorbed into enGenome’s commercial eVai platform. Commercial platforms, including VarSome (31) and Franklin (32), offer the most complete evidence, however, access to several key features is restricted by subscription paywalls. LLM-based tools are the newest variant curation assistant. VarChat (33) uses retrieval-augmented generation to summarise variant-specific literature and AI-CURA (34) separates automatable criteria from an LLM-supported assessment of literature-dependent evidence. AI-CURA is disease-agnostic and reported as a workflow rather than a deployed application, while the tools that are deployed and disease-tuned are proprietary and not open source.

**Table 1.**
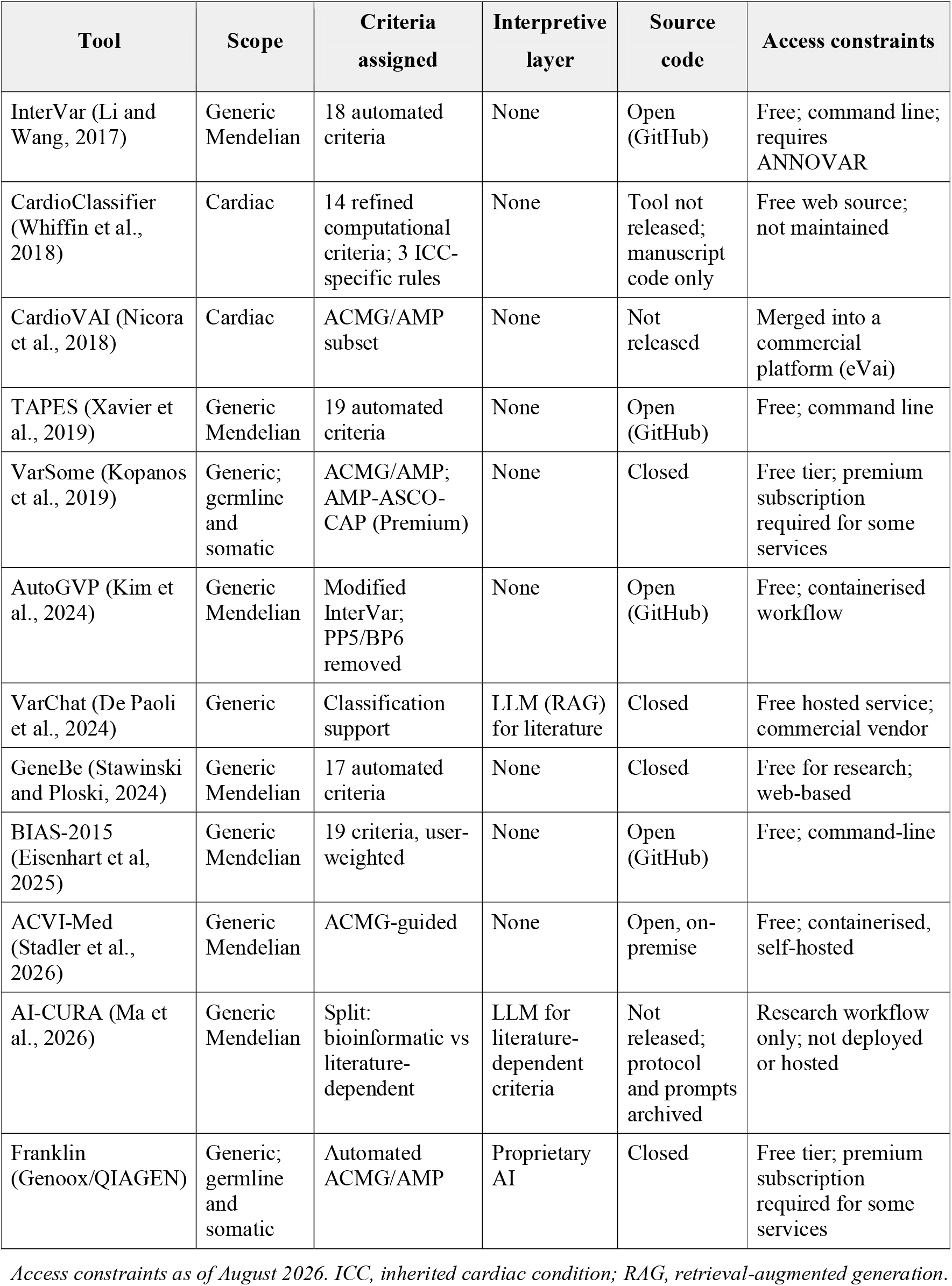
Software for ACMG/AMP variant classification.

| Tool | Scope | Criteria assigned | Interpretive layer | Source code | Access constraints |
| --- | --- | --- | --- | --- | --- |
| InterVar (Li and Wang, 2017) | Generic Mendelian | 18 automated criteria | None | Open (GitHub) | Free; command line; requires ANNOVAR |
| CardioClassifier (Whiffin et al., 2018) | Cardiac | 14 refined computational criteria; 3 ICC-specific rules | None | Tool not released; manuscript code only | Free web source; not maintained |
| CardioVAI (Nicora et al., 2018) | Cardiac | ACMG/AMP subset | None | Not released | Merged into a commercial platform (eVai) |
| TAPES (Xavier et al., 2019) | Generic Mendelian | 19 automated criteria | None | Open (GitHub) | Free; command line |
| VarSome (Kopanos et al., 2019) | Generic; germline and somatic | ACMG/AMP; AMP-ASCO-CAP (Premium) | None | Closed | Free tier; premium subscription required for some services |
| AutoGVP (Kim et al., 2024) | Generic Mendelian | Modified InterVar; PP5/BP6 removed | None | Open (GitHub) | Free; containerised workflow |
| VarChat (De Paoli et al., 2024) | Generic | Classification support | LLM (RAG) for literature | Closed | Free hosted service; commercial vendor |
| GeneBe (Stawinski and Ploski, 2024) | Generic Mendelian | 17 automated criteria | None | Closed | Free for research; web-based |
| BIAS-2015 (Eisenhart et al., 2025) | Generic Mendelian | 19 criteria, user-weighted | None | Open (GitHub) | Free; command-line |
| ACVI-Med (Stadler et al., 2026) | Generic Mendelian | ACMG-guided | None | Open, on-premise | Free; containerised, self-hosted |
| AI-CURA (Ma et al., 2026) | Generic Mendelian | Split: bioinformatic vs literature-dependent | LLM for literature-dependent criteria | Not released; protocol and prompts archived | Research workflow only; not deployed or hosted |
| Franklin (Genoox/QIAGEN) | Generic; germline and somatic | Automated ACMG/AMP | Proprietary AI | Closed | Free tier; premium subscription required for some services |

Access constraints as of August 2026. ICC, inherited cardiac condition; RAG, retrieval-augmented generation.

No currently available tool is simultaneously (i) parameterised for cardiovascular disease, (ii) able to evaluate the interpretive criteria rather than deferring them to the user, (iii) deployed as a curator-facing application that exports its evidence, and (iv) transparent enough that its criterion thresholds and model prompts can be inspected and corrected, and (v) available at no cost to the user. CardioClassifier (29) and CardioVAI (30) proved cardiovascular parameterisation was feasible, however neither tool was able to accommodate the continued evolution of the ACMG framework and its evidence. Since then, LLM-assisted development has substantially reduced the researcher hours needed to build and maintain these tools, narrowing the advantage that once made commercial platforms the only sustainable option.

We present HeartVar, a tool that implements a constrained-LLM approach for cardiovascular disease variant assessment. For each variant it gathers evidence in parallel from 20 public databases in seconds and passes it to the LLM as a single structured block, so that every assertion is grounded in live source data rather than model recall. HeartVar resolves rule-based criteria in code and reserves the LLM for those requiring expert judgement. Criteria are combined and the final tier is assigned programmatically. Every database annotation and criterion rationale is produced for manual interrogation, and the tool is explicitly designed to support curator review rather than autonomous reporting. By grounding every criterion in up-to-date evidence, HeartVar makes fast variant curation reproducible and auditable.

## Methods

### Inputs

A HeartVar query consists of a gene symbol and a variant (HGVS or genomic coordinates), together with optional proband information such as phenotype, inheritance, sex, and family history.

Both GRCh37 and GRCh38 coordinates can be used as input. GRCh37 coordinates are lifted to GRCh38 through Ensembl’s assembly mapping service before any other source is queried. A position returning no mapping or more than one aborts the curation with an explicit error rather than proceeding on an assumed coordinate. The curator’s original GRCh37 coordinates are retained and displayed alongside the lifted GRCh38 form.

### Evidence sources

HeartVar draws from the 20 sources summarised in Table 2. Each is used under its own licence and terms of use, and the primary publication for each source is cited. Most sources are served from local data files for speed, offline resilience, and to avoid rate-limiting under multi-user load. ClinVar (14), BioGRID (21), GTEx (19), UniProt (17), MGI (20), MedGen (35), fetal cell expression (36), and gnomAD (7) gene-constraint metrics are held in SQLite databases. gnomAD allele frequencies are served by tabix-slicing the gnomAD v4.1 sites VCFs, and SpliceAI (13) and AlphaMissense (12) are likewise tabix-indexed slices of their precomputed hg38 releases. Ensembl VEP (37) runs against a local cache. PanelApp (16), GenCC (15), Open Targets (38), CHDgene (39), and AlphaFold (40) are localised snapshots. ProtVar (18) annotations are assembled from the local AlphaMissense and UniProt records rather than queried remotely. Sources without a practical local mirror (PubMed, PubTator3 (41), and PMC Open Access) are queried live behind a shared TTL cache with single-flight de-duplication. Each source returns a structured response, including a deliberate error message when the source is not found, so the downstream prompt can distinguish absent from lookup failed. Data sources are updated monthly.

**Table 2.** HeartVar database sources.

| Source | Contribution |
| --- | --- |
| AlphaFold | Predicted 3D protein structure visualisation |
| AlphaMissense | Calibrated per-missense pathogenicity score |
| BioGRID | Curated protein-protein interactions; CHDgene partners flagged |
| CHDgene | Victor Chang Cardiac Research Institute curated congenital heart disease gene list |
| ClinVar | Clinical-significance reports, review status, star rating |
| Ensembl VEP | HGVS to GRCh38 coordinates, transcript consequence, SIFT, PolyPhen, REVEL, CADD |
| GenCC | Gene-disease classifications with submitter and mode of inheritance |
| gnomAD v4 | AF, popmax FAF95, hom counts, pLI/LOEUF/mis_z |
| GTEx v10 | Median TPM in left ventricle and atrial appendage |
| Heart of Fetal Cells | Fetal cardiac single-cell expression (36) |
| MedGen | NCBI gene-condition associations with OMIM-derived MIM numbers |
| MGI (via Alliance) | Mouse orthologue knockout/knockdown phenotypes; cardiac MP terms |
| Open Targets | Gene-disease association scores (genetic association, animal model, literature), HPO-matched where possible |
| PanelApp (Australia) | Cardiovascular panels, gene confidence, HPO match |
| PMC Open Access | Full-text Results/Methods/table excerpts for literature-gated criteria |
| ProtVar (EBI) | Residue-level functional and structural annotation |
| PubMed | Variant-specific and gene-level abstracts |
| PubTator3 | NCBI text-mining (tmVar3); recovers variant papers missed by strict PubMed queries |
| SpliceAI | Delta-acceptor/delta-donor splice impact |
| UniProt | Protein domains, active and binding sites, natural-variant annotations |

### ACMG/AMP framework

HeartVar applies the Richards et al. (2015) framework with the point-based combining rule of Tavtigian et al. (2020) (1, 2). Evaluation is split between a deterministic layer and the model. Nineteen of the criteria are computed programmatically in Python before anything reaches the LLM (BA1, BS1, BS2, BS4, PVS1, PS1, PS2, PM1, PM2, PM3, PM4, PM5, PM6, PP1, PP3, BP2, BP3, BP4, and BP7), because the evidence they rely on is structured and rule based. The model is asked to evaluate seven interpretive criteria that require human-level judgement (PS3, PS4, PP2, PP4, BS3, BP1, and BP5). It is noted that PP5 and BP6 are excluded from point-scoring due to updated recommendations regarding circularity of evidence (42).

### Model prompt and output schema

After annotating the variant, HeartVar assembles a prompt containing the clinical context, if provided, and the annotations and issues a single streaming call to the LLM. The model returns a single JSON object. It does not emit a classification or a point total. The server computes this from the model’s seven criteria plus the 19 precomputed criteria. The browser renders the result across the Summary, Variant, Gene, Protein, Criteria, and Export tabs.

### Large Language Model

This tool uses Anthropic Claude Sonnet 4.6 (temperature = 0) for AI interpretation. The use of AI is optional. Without AI, HeartVar still annotates every data source and returns a preliminary, rule-based ACMG/AMP classification computed from the criteria it can evaluate programmatically. The judgement-based criteria are marked “not assessed”, so there is no over-calling based on missing evidence. If the user opts to include AI interpretation (requires single sign-on), the full ACMG/AMP classification is returned as described above.

### Benchmarking

To evaluate classification performance, we assembled a benchmark of 106 cardiovascular disease variants from ClinGen carrying a three-star expert finding (Table S1) (43, 44). The variant set spanned several cardiovascular conditions (dilated cardiomyopathy, hypertrophic cardiomyopathy, long QT syndrome, RASopathy, and pulmonary hypertension) and covered 27 genes. Additionally, variants were distributed across ACMG tiers and included 22 Pathogenic, 18 Likely Pathogenic, 28 Variant of Uncertain Significance, 15 Likely Benign, and 23 Benign. The variants were submitted to HeartVar using the same information available to the original curator (gene, variant, phenotype, inheritance, zygosity, and *de novo* status), and the resulting ACMG/AMP tier and applied criteria were compared to the expert. Benchmarking was performed both with and without AI interpretation enabled, so that the contribution of the interpretive layer could be isolated. We report exact tier concordance and the direction and magnitude of any mismatch (over-versus under-calling). Performance was reported as per-criterion precision, recall, and F1 score, where precision was the proportion of tool-applied criteria also applied by the panel, recall was the proportion of panel-applied criteria also applied by the tool, and F1 was the harmonic mean of the two.

### Comparison with existing classifiers

HeartVar was compared against InterVar (23), GeneBe (28), and VarSome (31) on the same 106-variant set, each run in its default configuration with the same information available to the expert where possible. The remaining Table 1 tools were not benchmarked. AI-CURA (34) is released only as a workflow protocol rather than runnable software. CardioVAI (30) has been retired into enGenome’s commercial eVai platform. CardioClassifier (29) is operational but unsuited to our cohort because it is GRCh37 only, has not been updated since 2020, and limited to a gene panel that excludes several of the conditions assessed in our benchmark dataset. VarChat (33) returns its interpretation as free-text prose rather than a structured tier and criteria, so its output cannot be scored against the expert reference in the standardised way applied to the other tools. The rest offered little beyond the engines already included: TAPES (24) and AutoGVP (25) are InterVar-derivatives, while ACVI-Med (27) and BIAS-2015 (26) require multi-container or tens of gigabyte annotation backends disproportionate to their incremental value.

### Ethics

This study analysed only publicly available, de-identified variant interpretations from the ClinGen Evidence Repository. No identifiable patient information or human participants were involved, and therefore ethics committee approval and informed consent were not required.

## Results

HeartVar produces a preliminary ACMG/AMP classification with its full supporting evidence, intended to support expert variant assessment. To illustrate typical output, we describe classification of the *MYBPC3* NM_000256.3:c.1483C>G variant in a proband with autosomal dominant hypertrophic cardiomyopathy (Figure 1). On submission, evidence is aggregated in parallel from 20 sources, and ACMG/AMP criteria are evaluated before being scored under the Tavtigian et al. (2020) point system (2). For this particular variant, programmatic evaluation applied PM1_Moderate (+2), PM2_Supporting (+1), and PP1_Strong (+4). The AI interpretation layer additionally supported PS4_Supporting (+1) sourced from relevant literature. Summed under the Tavtigian point system, the variant reaches +8 points, yielding a draft classification of Likely Pathogenic. This classification and four criteria match the evidence in the ClinGen eRepo expert-panel record. The interface presents each criterion and its supporting evidence alongside the aggregated source data, so the curator can trace the classification to its underlying evidence.

**Figure 1.**
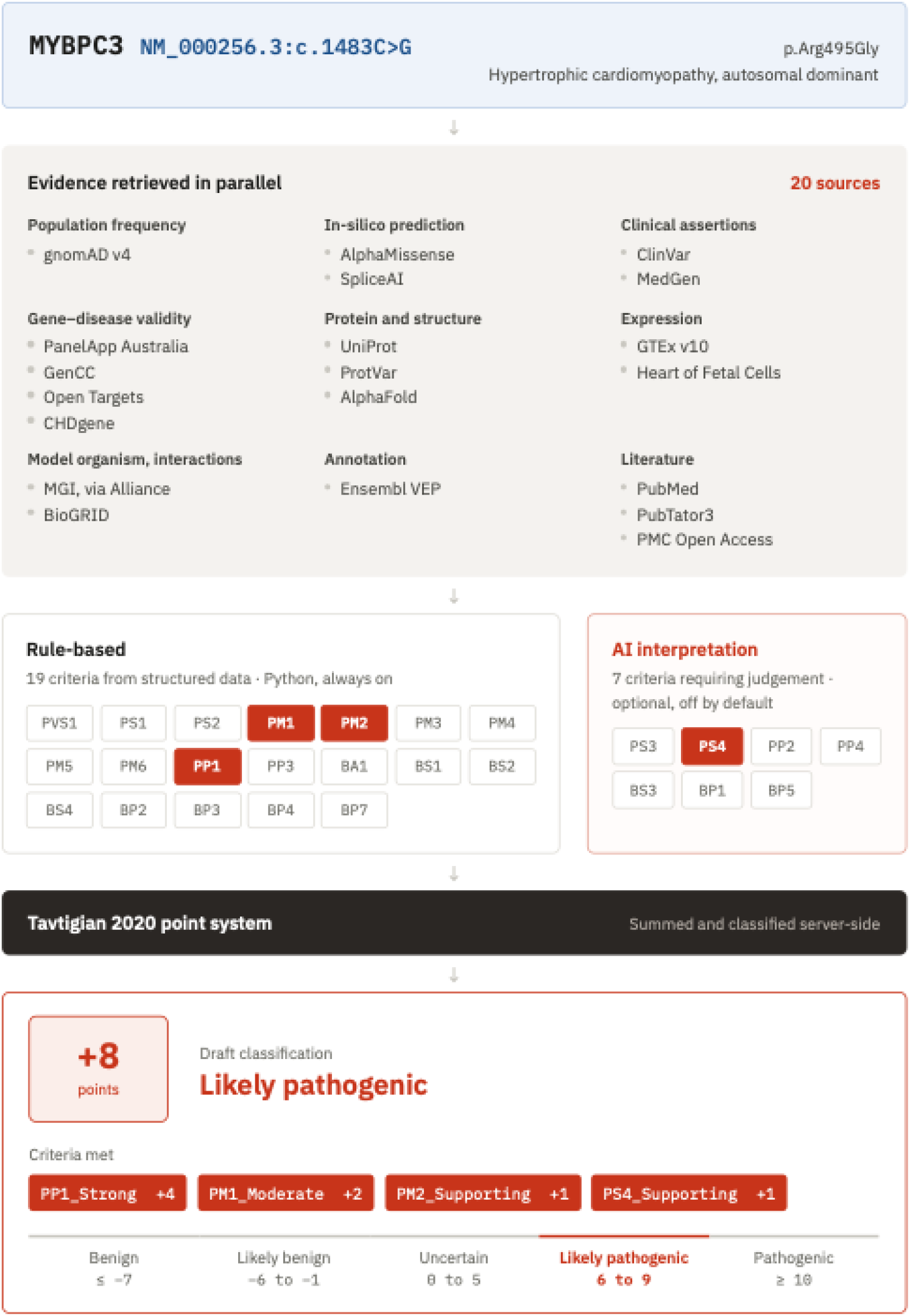
Overview of the HeartVar variant curation pipeline. For the worked example *MYBPC3* NM_000256.3:c.1483C>G in hypertrophic cardiomyopathy, evidence is aggregated in parallel from 20 sources and used to evaluate ACMG/AMP criteria: 19 rule-based and seven via an optional LLM call. Met criteria are summed under the Tavtigian (2020) point system to give a preliminary, expert-reviewable classification (here +8 points, Likely Pathogenic).

### Benchmark performance

Across the 106 variant benchmark set, HeartVar reproduced the reference ACMG/AMP tier in 72% of variants with per-criterion precision of 72% (Table 3). The confusion matrix (Table 4) shows that errors were largely conservative and, with one exception, mismatches were limited to the adjacent tier. Of the 22 expert P variants, HeartVar called 11 P and 7 LP, with the remaining 4 under-called to VUS; of the 18 expert LP variants, 13 were concordant, 4 were under-called to VUS, and 1 over-called to P. No expert P/LP variant was called LB/B, and *vice versa*. On the benign side agreement was stronger, with 21 of 23 expert B and 9 of 15 expert LB variants matched exactly (Table S2-3).

**Table 3.**
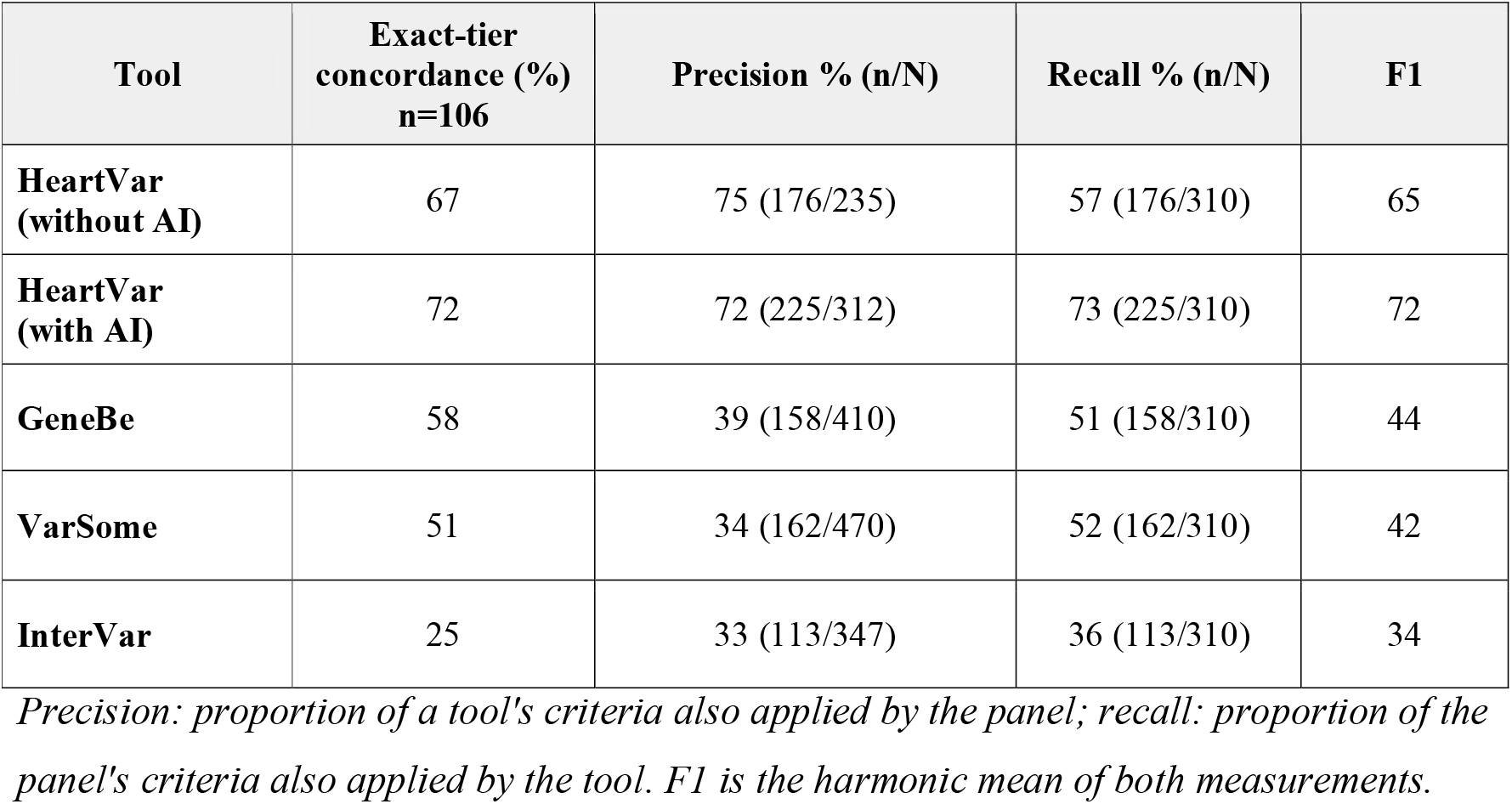
Benchmarking results for cardiovascular disease variants from ClinGen eRepo for HeartVar and other variant curation tools. (n = 106 variants; 27 genes; P = 22, LP = 18, VUS = 28, LB = 15, B = 23).

| **Tool** | **Exact-tier concordance (%)**<br>**n=106** | **Precision % (n/N)** | **Recall % (n/N)** | **F1** |
| --- | --- | --- | --- | --- |
| **HeartVar (without AI)** | 67 | 75 (176/235) | 57 (176/310) | 65 |
| **HeartVar (with AI)** | 72 | 72 (225/312) | 73 (225/310) | 72 |
| **GeneBe** | 58 | 39 (158/410) | 51 (158/310) | 44 |
| **VarSome** | 51 | 34 (162/470) | 52 (162/310) | 42 |
| **InterVar** | 25 | 33 (113/347) | 36 (113/310) | 34 |

*Precision: proportion of a tool's criteria also applied by the panel; recall: proportion of the panel's criteria also applied by the tool. F1 is the harmonic mean of both measurements.*

**Table 4.** HeartVar benchmarking confusion matrix. Results from evaluation using full proband context and LLM interpretive layer (n = 106 variants).

|  | **Pathogenic**<br>**(HV)** | **Likely**<br>**pathogenic**<br>**(HV)** | **Variant of**<br>**uncertain**<br>**significance**<br>**(HV)** | **Likely**<br>**benign**<br>**(HV)** | **Benign**<br>**(HV)** | **Total** |
| --- | --- | --- | --- | --- | --- | --- |
| **Pathogenic**<br>**(E)** | 11 | 7 | 4 | 0 | 0 | 22 |
| **Likely**<br>**Pathogenic**<br>**(E)** | 1 | 13 | 4 | 0 | 0 | 18 |
| **Variant of**<br>**uncertain**<br>**significance**<br>**(E)** | 0 | 2 | 22 | 4 | 0 | 28 |
| **Likely**<br>**benign (E)** | 0 | 0 | 6 | 9 | 0 | 15 |
| **Benign (E)** | 0 | 0 | 1 | 1 | 21 | 23 |
| **Total** | 12 | 22 | 37 | 14 | 21 | 106 |

*E = expert variant curation; HV = HeartVar curation*

### Contribution of the LLM interpretive layer

Isolating the interpretive layer by benchmarking HeartVar with and without AI interpretation showed that the LLM improved exact-tier concordance and strengthened recovery of expert-applied criteria (Table 3; Table S2-3). Adding the AI interpretation raised exact-tier concordance from 67% to 72%. Per-criterion precision fell slightly (75% to 72%) while recall rose sharply (57% to 73%), lifting F1 from 65% to 72%, demonstrating the value of the LLM layer for interpretive criteria.

### Comparison with existing classifiers

HeartVar performed as well as or better than existing classifiers. The ACMG/AMP tier matched the expert classification for 72% of variants with HeartVar, compared with 58% for GeneBe, 51% for VarSome, and 25% for InterVar (Table 3; Figure 2; Table S3-6). The tools differed not only in accuracy, but in the direction and basis of their errors. HeartVar’s discordances were largely conservative. Of the 40 expert pathogenic (P/LP) variants, 32 were called P/LP and the remainder were under-called to VUS. VarSome erred in the opposite direction, systematically over-calling by a tier (Figure 3). It returned 17 of 18 LP variants as P and reassigned 16 of 28 expert VUS variants to LP/P (Table S4). GeneBe’s concordance rested heavily on ClinVar assertions rather than independent evidence evaluation. From exact matches, 47 out of 61 (77%) applied PP5 or BP6. Every one of its correct P/LP calls applied PP5, while every correct LB/B call applied BP6 (Table S5).

**Figure 2.**
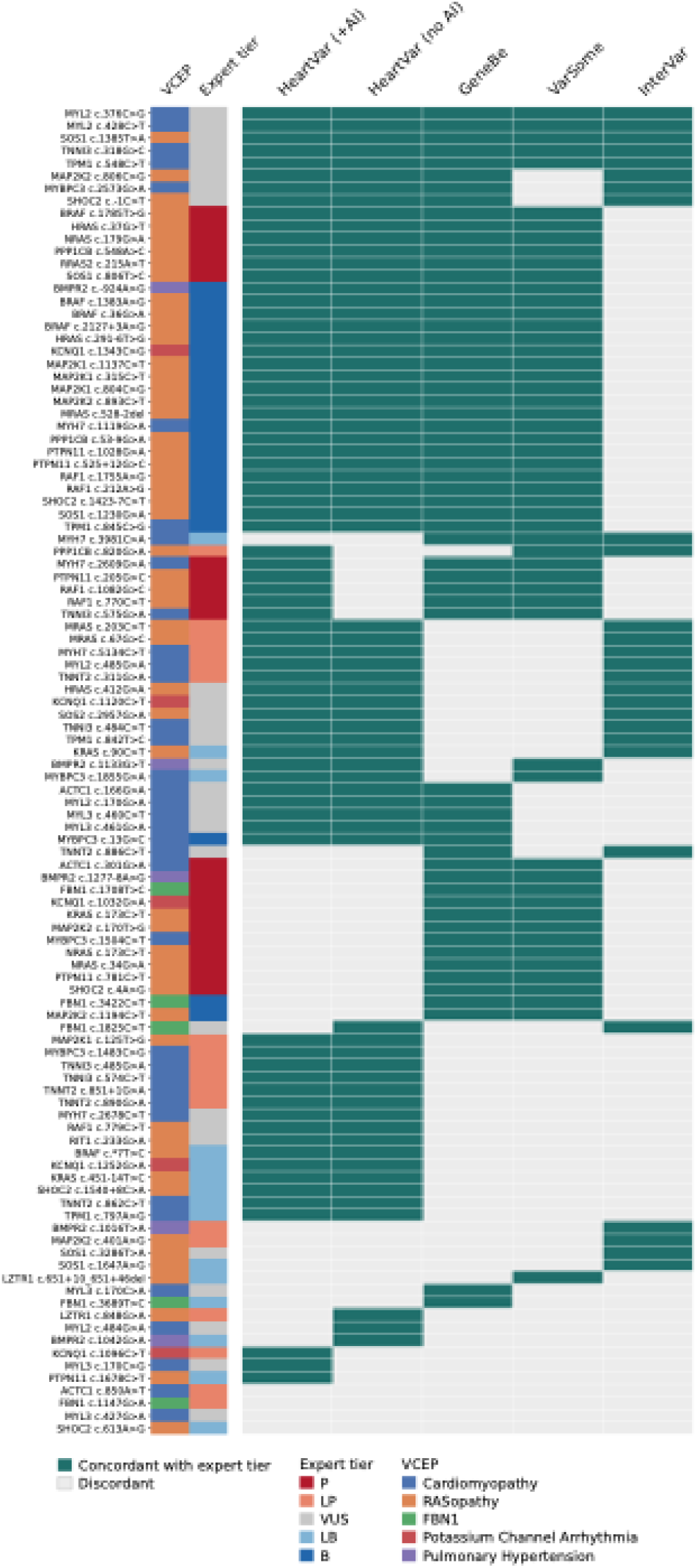
Variant-level concordance with ClinGen expert classifications. Heatmap showing whether each classification tool assigned the same ACMG/AMP tier as the expert panel for 106 benchmark variants. Teal indicates concordance and grey discordance. Rows are ordered by shared concordance pattern and columns by overall concordance; side annotations show the VCEP category and expert tier. P, pathogenic; LP, likely pathogenic; VUS, variant of uncertain significance; LB, likely benign; B, benign.

**Figure 3.**
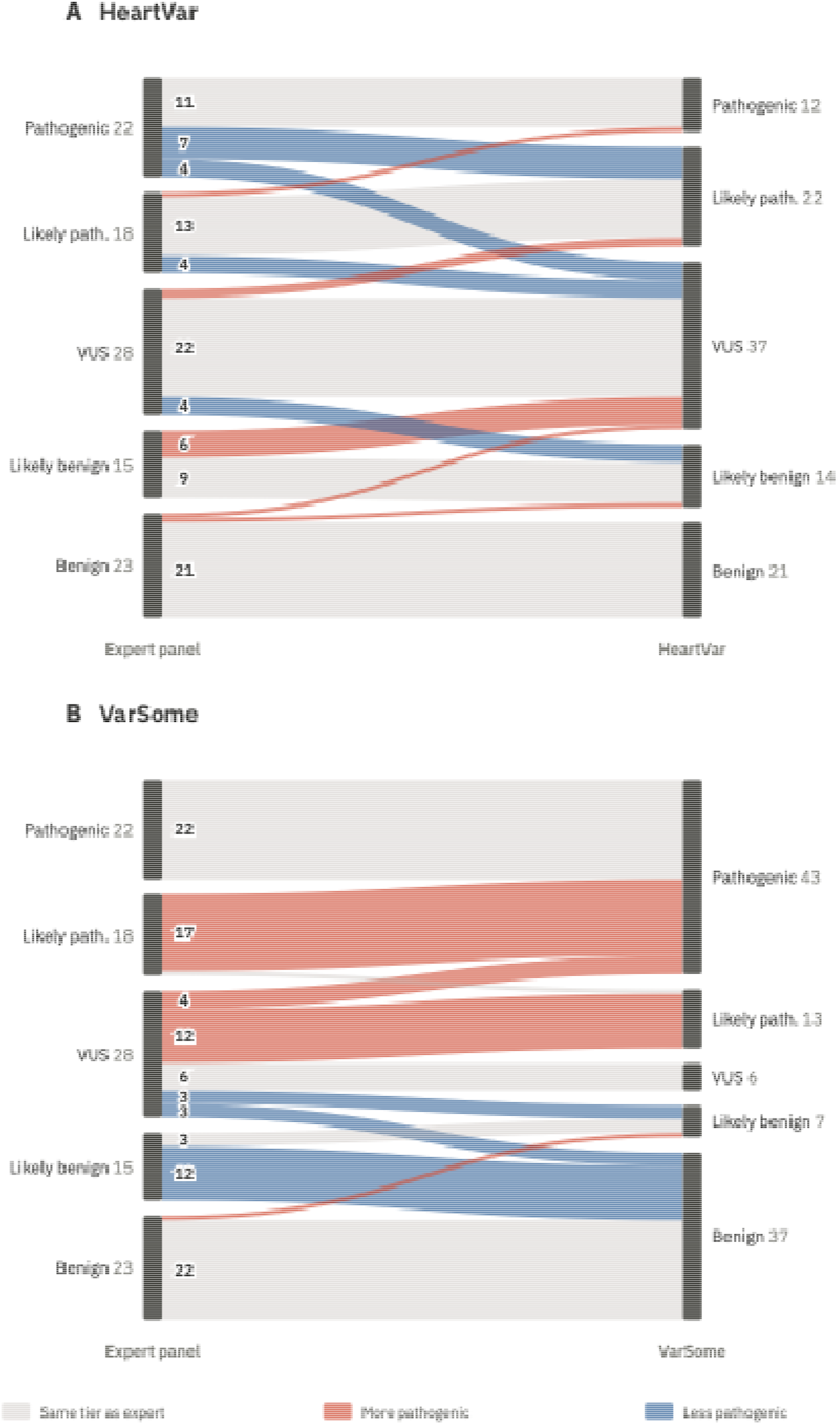
Comparison of A) HeartVar and B) VarSome variant classifications against expert-panel variant interpretations. Sankey diagrams compare ACMG/AMP tiers assigned to 106 ClinGen expert-curated cardiovascular disease variants by expert panels, HeartVar, and VarSome. Band widths represent the number of variants in each classification flow. P: pathogenic; LP: likely pathogenic; VUS: variant of uncertain significance; LB: likely benign; B: benign.

## Discussion

HeartVar is the first cardiovascular disease-specific ACMG/AMP tool that is freely available, open source, and deployed as a curator-facing application. It combines disease-specific parameterisation, evaluation of interpretive criteria, exportable per-criterion evidence, and transparent prompts. The LLM layer improves performance, particularly by recovering expert-applied criteria that depend on interpretation of unstructured evidence. It also reduces the manual effort required to assess evidence distributed across primary literature, including functional studies, case enrichment, segregation, and phenotype specificity. This changes the nature of curator review rather than removing the need for it. Because LLMs can misread or fabricate evidence, each model-assigned criterion is shown with the source data on which it was based, allowing curators to verify the proposed interpretation. Grounding the model in 20 specific data sources rather than model recall keeps these judgements traceable, while assigning the final tier by rule rather than by the model keeps the classification reproducible.

Maintenance is also central to the usefulness of variant curation tools. A major challenge for academic variant curation tools has been sustainability: evidence sources, ACMG/AMP specifications, gene-disease validity assertions, and software dependencies all change over time. LLM-assisted development will reduce some of this maintenance burden by lowering the researcher time required to update data sources, revise criteria, and keep the tool aligned with evolving best practices.

The benchmark results should be read with consideration of HeartVar’s intended role. Its disagreements with expert panels were largely conservative under-calls, reflecting case and functional evidence available to expert curators that is not fully recoverable from public literature. This conservative bias is the safer failure mode for a first-pass assistant: it avoids over-calling pathogenicity while still surfacing evidence for curator assessment and, where warranted, tier reassignment. HeartVar is therefore positioned to accelerate expert curation, not replace it. Its per-criterion transparency allows curators to accept, adjust, or overturn any assignment against the evidence shown.

Two variants classified as VUS by the expert panel were classified as Likely Pathogenic by HeartVar. For *FBN1* c.1825C>T, HeartVar applied PP2 (missense variant in a gene with a low rate of benign missense variation and where missense variants are a common mechanism of disease) and PP4 (patient phenotype or family history highly specific for a disease with a single genetic aetiology), both of which the panel recorded as not met, and did not apply BS1 (allele frequency greater than expected for the disorder). However, the allele frequency supporting BS1 in the expert review was present in gnomAD v2.1.1 but falls below the *FBN1* VCEP threshold in gnomAD v4.1, indicating that this benign evidence is absent from the newer dataset rather than missed by HeartVar. For *MYL2* c.484G>A, HeartVar matched the panel’s reduced PM2 (absent from controls) and PM5 (novel missense change at an amino acid residue where a different pathogenic missense change has been seen before) strengths, but applied PM6 (assumed *de novo*) at Moderate instead of Supporting and applied PS3 (well-established functional studies demonstrate a deleterious effect on the gene) based on published overexpression and model-organism assays the panel did not credit. Conversely, HeartVar missed the panel’s PS4 (prevalence of the variant in affected individuals is significantly increased compared with controls), which was based on odds-ratio evidence that could not be derived from the open-source literature retrieved by the tool. These discrepancies reflect the expert panel’s access to case-level and phenotypic detail unavailable to HeartVar, reinforcing the need for manual review.

The benchmarking tools showed distinct error profiles. VarSome’s systematic upward shift inflates apparent pathogenic sensitivity at the cost of specificity, the opposite of HeartVar’s conservative bias. GeneBe’s concordance rests largely on PP5/BP6 (a reputable source reports the variant as pathogenic/benign, without supporting evidence), which assign evidence from an existing reputable-source classification. Current recommendations are that both criteria be discontinued because they reintroduce prior classifications as evidence and risk circularity (42). Because the benchmark variants were drawn from the ClinGen Evidence Repository, GeneBe’s use of these criteria effectively reads the reference classification back rather than reconstructing it from primary data; its accuracy would therefore be expected to fall for variants without an existing ClinVar assertion. HeartVar follows current recommendations by excluding PP5 and BP6 from scoring while still surfacing that information for curator review, so its agreement with expert calls is built from primary evidence rather than prior classifications.

Several limitations should be considered when using HeartVar. First, it is parameterised and validated only for cardiovascular disease, so its performance in other domains is unknown. The gene panel, ClinGen thresholds, criterion-applicability rules, expression resources, and phenotype matching are all cardiovascular-specific. Source code is publicly available, however, and the tool can be adapted to other disease areas. Second, the ClinGen eRepo variants used for benchmarking come from existing expert assertions and may not represent novel or conflicting variants, where first-pass support may be most valuable. The benchmark also does not fully represent all cardiovascular disease, and performance may vary across less well-represented subdomains. Finally, the interpretive layer depends on a commercial LLM, introducing versioning and reproducibility considerations. This is mitigated by rule-based tier assignment and by showing every model assertion for curator review.

HeartVar is a cardiovascular-focused, LLM-assisted variant curator that searches 20 evidence sources, applies the current ACMG/AMP framework, and surfaces each criterion assignment for curator review. As an open-source tool, it provides a reproducible starting point for clinical genomics laboratories running cardiovascular studies and for methods researchers developing LLM-assisted approaches to variant interpretation.

## Acknowledgements

We thank the maintainers of the public resources HeartVar builds on: Ensembl/EMBL-EBI, the Broad Institute (gnomAD), Illumina (SpliceAI), Google DeepMind (AlphaMissense, AlphaFold), NCBI (ClinVar, PubMed, PubTator3, MedGen), the UniProt Consortium, EBI ProtVar, the GTEx Consortium, PanelApp Australia, the Gene Curation Coalition (GenCC), the Alliance of Genome Resources/MGI, BioGRID, Open Targets, and Farah et al. for the Heart of Fetal Cells dataset. CHDgene is maintained by the Victor Chang Cardiac Research Institute.

## Data availability

HeartVar is hosted free of charge at www.heartvar.victorchang.edu.au. Source code is available at github.com/VCCRI/heartvar-open under the MIT licence.

## AI Disclosure

HeartVar was developed with extensive use of AI-assisted programming (Anthropic Claude models, accessed via the Anthropic API and Claude Code). Anthropic’s Claude was additionally used to assist with drafting and editing prose, structuring sections, and reviewing analysis and figure generation. All AI-assisted content was reviewed and revised by the authors, who designed the study, ran and interpreted the analyses, and produced the final tables and figures. The authors take full responsibility for the content of the publication.

